# Antagonistic activity of AEA on TAT treated Human Astrocytes identify Inflammaging pathways

**DOI:** 10.1101/2024.10.21.619556

**Authors:** Durairaj Duraikkannu, Kamini Khatak, Hemavathy Nagarajan, Sridharan Sudharshan, Nivedita Chatterjee

**Affiliations:** L&T Department of Ocular Pathology, Vision Research Foundation, 41 College Road, Chennai, India 600006; Centre for Biotechnology, Anna University, Chennai 600025, India, Research Scholar; Department of Bioinformatics, Vision Research Foundation, 41 College Road, Chennai, India 600006; Department of Uveitis, Medical Research Foundation, 41 College Road, Chennai, India 600006; Ion Channel Biology Laboratory, AU-KBC Research Centre, Madras Institute of Technology, Anna University, Chennai-600 044, Tamil Nadu, India

**Keywords:** Endocannabinoid, AEA, HIV, senescence, Inflammaging

## Abstract

Physiologically, the endocannabinoids are known to reduce inflammation by decreasing production of inflammatory factors in immune and glial cells. Astrocytes secrete soluble inflammatory mediators and prolonged activation of astrocytes is associated with accelerated aging in Central Nervous System. Many reports show miRNAs as critical gene regulators in inflammation and astrogliosis in astrocytes. The aim of this study is to investigate the microRNA changes affected by Anandamide on HIV1 TAT (Trans-activator of transcription protein) stimulated normal human astrocytes. We performed global microRNA profile in TAT activated astrocytes and analysed changes on exposure to AEA. To delineate the mechanism of action we assessed with bioinformatic tools miRWalk, KEGG and Cytoscape the global microarray for significantly impacted miRNAs, and their gene targets. TAT activation in astrocytes upregulated 122 miRNAs significantly (p < 0.05). Addition of AEA in activated astrocytes downregulate the expression of 57 miRNAs significantly. Out of 122 upregulated miRNAs on TAT treatment, 37 miRNAs which were common in TAT and TAT+AEA cells showed reversal suggesting clues to critical miRNAs for the AEA-induced mitigation of neuroinflammation. Reversal in expression of selected group of miRNAs identify antagonistic pathways which are promoting anti-inflammatory environment. Pathway analysis of key 37 miRNAs show gene targets that regulate inflammation and senescence.

## Introduction

The endogenous cannabinoid system modulates immune phenomena associated with infection or inflammation. Due to their ability to suppress lymphocyte proliferation and inflammatory cytokine production, there is interest in them as promising therapeutic targets for diseases with immune phenotype (Lowe et al. 2021). Because all immune cells examined so far express cannabinoid receptors regardless of their cell lineage, all types of immunity are sensitive to cannabinoid modulation. Studies on SIV infected macaques upon ΔTHC administration and investigations on people living with HIV using cannabis has also shown decreased levels of T-cell activation, inflammatory monocytes and pro-inflammatory cytokine secretion (Mehraj et al.2019). Endocannabinoids (EC) can modulate immune reactions in the CNS through their effects on microglia (Tanaka et al. 2020), astrocytes [4], and oligodendrocytes (Ilyasov et al. 2018). Two endocannabinoids, Anandamide (AEA) and 2-Arachidonoylglycerol (2-AG) affect innate immune response of the retinal glia Muller cells (Krishnan & Chatterjee et al. 2012, 2014, 2015) The authors had earlier shown AEA and 2-AG work to reduce inflammation by critical targeting of inflammation machinery at multiple signalling levels.

In CNS, despite viral suppression by anti-retroviral therapy (ART), the symptoms of HIV-associated neurocognitive disorder (HAND) endure. Instead of resident T-lymphocytes, the virus in the brain lives on in microglia and astrocytes, establishing a reservoir of infection. The HIV proteins, some secreted such as HIV1 TAT (TAT) can inflame neurons causing neurodegeneration. The HIV1 TAT protein is a regulatory protein that highly enhances transcription. Persistent chronic inflammation underlying molecular mechanisms lead to premature aging. Triggered astrocytes secrete soluble mediators, such as CXCL10, CCL2, interleukin-6 and BAFF, causing glial dysfunction and contributing to HIV-1 neuropathogenesis. The role of miRNAs in regulating the aging, inflammation process has also been well studied (Doke et al. 2021). These small non-coding RNAs (21 – 25 nucleotides long) regulate the gene expression by degrading or silencing their target mRNA molecules by using their complementary seed sequences. miRNAs bind with the 3’UTR (Untranslated region) or 5’UTR of their target mRNAs. Complexities of interactions are increased by numerous binding sites per miRNA and the potential of each mRNA to be targeted by multiple miRNAs (Krol et al. 2010, O’neill et al. 2011) . In this study we investigate the effects of AEA on HIV1 TAT activated cultured human astrocytes. We look at the global changes in microRNAs by AEA, which are involved in the regulation of inflammation particularly as it relates to inflammaging. The endocannabinoid AEA has already been shown to downregulate key miRNAs that attenuate inflammation in a Staphylococcus Enterotoxin B (SEB) model of acute respiratory inflammation model through down-regulating immunosuppressive T regulatory cells (Sultan et al. 2021)

In a global miRNA microarray analysis, addition of TAT to astrocytes significantly increases the expression of 122 miRNAs. We analysed the anti-inflammatory role of AEA in the TAT stimulated astrocytes. AEA addition along with TAT, show decrease in majority of miRNAs, many related to inflammation. Thirty seven miRNAs (out of 122 miRNAs in TAT treated cells) which were common to both treatments showed reversal suggesting their role in systematic suppression and may yet provide suitable therapeutic targets by pharmacological intervention.

## Materials and methods

### Primary cell culture and treatment

Normal Human Astrocyte (Cat# CC-2565, Lonza) had been cultured with AGM^TM^ Astrocyte Growth Medium Bullet Kit^TM^ (basal medium and Single Quotes^TM^ Kit; Cat# CC-3186) for cell specific growth. Cells were seeded into a T75 culture flask and allowed to grow in CO_2_ incubator for a week to attain 80% confluency. Cells were split in 6 well plates for treatment. Once the cells attained 80% confluency, they were treated with HIV-1 TAT recombinant protein, AEA and TAT+AEA for 24 hours, and collected for microarray. TAT recombinant protein was purchased from NIH AIDS (ARP 2222). The lyophilized powder was reconstituted by using sterile PBS reagent. 100 ng/mL of TAT was used for treatment. AEA (Sigma) was reconstituted by using DMSO to attain the working concentration of 10 μM.

### RNA isolation

Treated cells were harvested by using TRIzol™ Reagent (Cat. No: 15596018) Invitrogen^TM^. By using chloroform reagent, phase separation was obtained. The clear upper aqueous phase was collected and the RNA was precipitated by using isopropanol. Isopropanol is removed then by using 70% ethanol. The precipitate was air dried and dissolved in RNase free water.

### Global microarray profiling

Global miRNA microarray profiling was done for human miRNAs by using Agilent’s complete miRNA labeling and Hybridization Kit in Human 8X60k AMADID: 70156 slides (Cat # 5190-0456). Expression of miRNAs was given in log 2-fold change. The cut off value was 0.6. miRNA expression of TAT and AEA treated samples were compared and normalized with untreated astrocyte (control). miRNA expression of TAT+AEA treated samples were normalized with TAT treated astrocytes. The normalization of miRNA expression was done using GeneSpring GX Software.

Normalize to specific samples: Treated vs. Control Percentile Shift Normalization is a global normalization, where the locations of all the spot intensities in an array are adjusted. This normalization takes each column in an experiment independently, and computes the percentile of the expression values for this array across all spots (where n has a range from 0-100 and n=50 is the median). It subtracts this value from the expression value of each entity.

### miRNA-mRNA Network

In our study, addition of TAT increases the expression of 122 miRNAs. When AEA and TAT are added together, expression of 37 (out of 122 miRNAs) miRNAs were brought down. These differentially expressed miRNAs were further analysed. The differentially expressed miRNAs were subjected to miRWalk database (http://mirwalk.umm.uni-heidelberg.de/) to predict their target mRNAs. Functional mRNAs that are having high binding energy (cut off set as 1) were shortlisted. Cytoscape software (https://cytoscape.org), was used to understand interactions between the shortlisted miRNAs and their predicted mRNAs.

### Pathway enrichment analysis

Shortlisted mRNAs were fed into KOBAS (KEGG Orthology-Based Annotation System) (https://bio.tools › kobas) pathway analysis tool to predict the pathways regulated by the miRNAs. By keeping the adjusted p value <0.05, significant pathways that are related to inflammation and aging were selected.

### Protein-protein interaction (PPI) network analysis

The Search Tool for the Retrieval of Interacting Genes (STRING) (http://string-db.org/) demonstrated the interactions between the proteins that are predicted as the targets for the shortlisted miRNAs.

### Comparative analysis of the target miRNAs

A comparative analysis was carried out for the resultant miRNA-mRNA with the known mRNAs of genes that are involved in inflammasome, senescence and endocannabinoid system, in order to validate with the miRNAs experimentally observed by us. The resultant mRNAs were subjected into miRDB (MicroRNA Target Prediction Database) (http://mirdb.org) server to predict their regulatory miRNAs. Subsequently, the common miRNAs in both miRDB predicted and our microarray profile were reported.

## Results

### miRNA profiling

HIV-1 TAT regulate transcription of host genes and miRNA through a number of transcription factors including CREB (Rahimian et al. 2016). miRNA expression is mainly regulated at the level of transcription. miRNA expression on TAT and TAT+AEA treatment is represented as heat map (Figure 1a) using GeneSpring GX software. Astrocytes on treatment with TAT protein upregulates 470 miRNAs. Out of these 470 miRNAs, 122 miRNAs showed statistically significant values (p < 0.05) (Figure 1b-c and Figure 2a). The fold change values of upregulated miRNAs are between 0.5 and 5.37 (hsa-miR-8485, hsa-miR-513a-5p, hsa-miR-96-5p, hsa-miR-543, hsa-miR-1271-5p are highly upregulated). Significant upregulation of 122 miRNAs indicate regulation of expression of many mRNAs which are controlled by these miRNAs. In our data, TAT downregulates two miRNAs (hsa-miR-4701-5p, hsa-miR-6763-3p).

**Figure 1.**
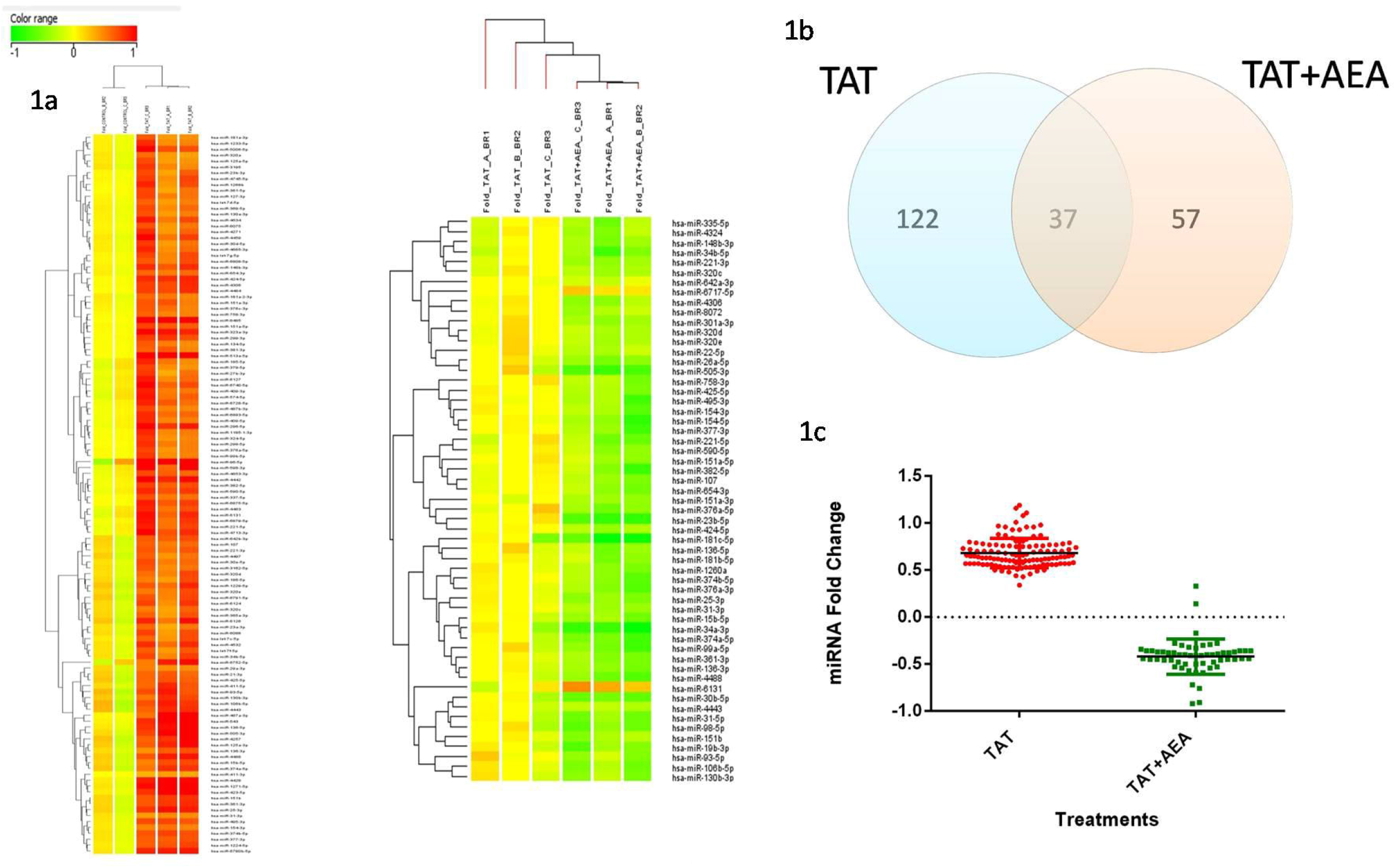
(a) AEA alters miRNA expression in human astrocytes treated with HIV1 TAT coat protein. Primary human astrocytes were screened for microRNA expression as described in Methods. Heat map show altered miRNA on TAT and TAT+AEA treatment. Red denotes upregulation and green denotes downregulation. Fold change range taken between -1 and 1. (b) TAT treatment in astrocytes increases the expression of 122 miRNAs significantly. TAT+AEA decreases the expression of 57 miRNAs. Out of 122 upregulated miRNAs 37 miRNAs are downregulated by AEA. (c) miRNA expression is represented as fold change log base 2 in TAT and TAT+AEA cells. AEA decreases expression of 37 miRNAs which were upregulated on TAT treatment.

**Figure 2.**
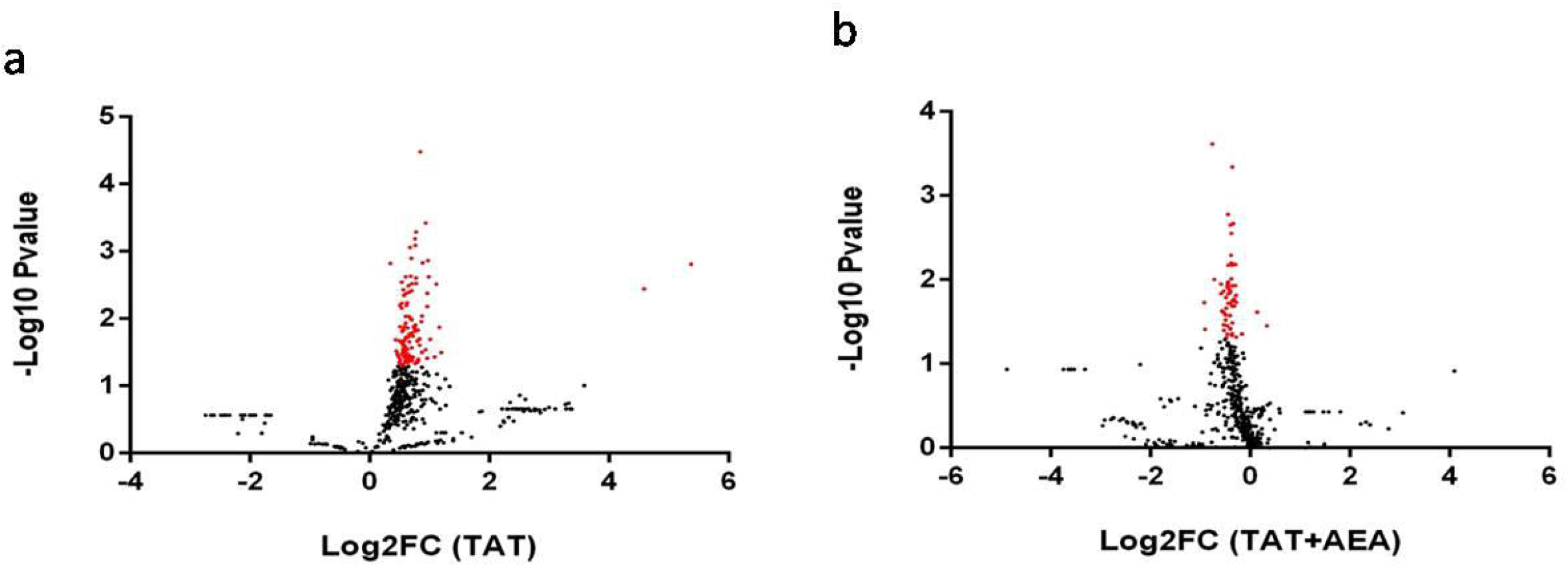
(a) Volcano plot representing the expression of miRNAs in TAT treated astrocytes. Majority of miRNAs are upregulated with TAT. miRNAs that are significantly upregulated appear in red. (b) Volcano plot representing the expression of miRNAs in TAT+AEA treated astrocytes. Majority of miRNAs are downregulated with TAT+AEA. miRNAs that are significantly downregulated appear in red.

Exposure to AEA increases the expression of 266 miRNAs when compared to untreated control. Out of 266 miRNAs, 52 miRNAs are significantly upregulated. hsa-miR-4788, hsa-miR-8485, hsa-miR-4698 (Doke et al. 2021), hsa-miR-6500-5p, hsa-miR-6777-3p are statistically significant. AEA treatment lead to one downregulated miRNA (hsa-miR-145-3p).

Fifty seven downregulated miRNAs show statistical significance out of the 386 miRNAs which change on treatment with TAT+AEA (Figure 2b), when compared to cells with TAT. Downregulation of 57 miRNAs indicate anandamide’s role in reversal of multiple regulatory elements. It is to be noted that the miRNAs hsa-miR-8485, hsa-miR-513a-5p, hsa-miR-96-5p, hsa-miR-543, hsa-miR-1271-5p which were upregulated by TAT were not downregulated by TAT+AEA. TAT+AEA treated astrocytes show upregulation in 7 miRNAs, none statistically significant (hsa-miR-6768-5p, hsa-miR-6780a-5p, hsa-miR-6880-3p, hsa-miR-2116-3p, hsa-miR-6512-5p, hsa-miR-3125, hsa-miR-6777-3p). Thus, AEA has an antagonistic role in TAT activated cells but not in resting cells. We have taken the statistically significant miRNAs for further analysis. Since we wanted to investigate the effect of AEA with TAT, we focused mainly on miRNA expression in TAT+AEA treated cells. Out of 57 miRNAs downregulated in TAT+AEA treatment, 37 miRNAs are common in both TAT (122 upregulated miRNAs) and TAT+AEA treatment. AEA therefore reverses the expression of large number of miRNAs upregulated by TAT. KEGG enrichment analysis was done for the target genes related to these 57 miRNAs. These miRNAs are related to PI3-Akt signaling pathway, Ras signaling pathway, MAPK signaling pathway, longevity regulating pathway, TNF signaling pathway and pathways related to cancer.

### Target prediction

We performed bioinformatic analysis of 37 miRNAs which were upregulated on TAT treatment and down regulated by AEA in TAT activated cells. Therefore, these 37 miRNAs hold a significant role in the antagonistic activity of AEA on TAT. The total numbers of target mRNAs predicted for each miRNA is given in the Supplementary Table 1 (high binding energy cut off set as 1). mRNAs that are regulated by more than one miRNA (Table 1 and 2) were only considered to generate the miRNA-mRNA networks. We found that miR-15b-5p, miR-93-5p and miR-106b-5p are predicted to regulate many genes that are associated with inflammation and senescence. miR-15b-5p, miR-93-5p and miR-106b-5p are upregulated to the fold change of 0.63, 0.76 and 0.79, respectively with TAT. On TAT+AEA addition these are downregulated to fold change of -0.44, 0.36 and -0.45, respectively. Significant predicted mRNAs include FOXO1, FOXO3, MAPK1, MAPK9, JAK1 FRS2, RPS6KA5 and RPS6KA1 as part of inflammation machinery and their miRNA-mRNA network image is given in Figure 3a. Predicted significant mRNAs related to senescence pathways identified are FOXO1, FOXO3, NR3C1, ATXN3, CAPZA2, AKT3, BCL2L11, TP53, TXNIP, CYCS, MAPK9, AGO3, DCTN5, RB1, CBX8, TGFBR2, MAPK1, PTEN, YWHAQ, AKT2 and RPS6KA1. Their miRNA-mRNA network image is given Figure 3b.

**Table 1.** From KEGG pathway enrichment analysis mRNAs are predicted. mRNAs that are regulated by more than one shortlisted miRNAs for inflammasome genes are given in the table.

|  | **miRNAs** | **Predicted targets (mRNAs)** |
| --- | --- | --- |
| **1** | **miR-15b-5p, miR-221-3p** | **FOXO1, FOXO3** |
| **2** | **miR-6131, miR-34b-5p, miR-93-5p, miR-130b-3p** | **NR3C1** |
| **3** | **miR-6131, miR-382-5p** | **ATXN3** |
| **4** | **miR-15b-5p, miR-93-5p** | **CAPZA2** |
| **5** | **miR-15b-5p, miR-221-3p** | **AKT3** |
| **6** | **miR-221-3p, miR-25-3p, miR-93-5p, miR-106b-5p** | **BCL2L11** |
| **7** | **miR-151a-3p, miR-25-3p, miR-221-3p** | **TP53** |
| **8** | **miR-31-3p, miR-130b-3p** | **TXNIP** |
| **9** | **miR-93-5p, miR-106b-5p** | **CYCS** |
| **10** | **miR-93-5p, miR-106b-5p** | **MAPK9** |
| **11** | **miR-93-5p, miR-106b-5p, miR-130b-3p** | **AGO3** |
| **12** | **miR-93-5p, miR-106b-5p** | **DCTN5** |
| **13** | **miR-93-5p, miR-130b-3p** | **RB1** |
| **14** | **miR-93-5p, miR-106b-5p** | **CBX8** |
| **15** | **miR-106b-5p, miR-130b-3p** | **TGFBR2** |
| **16** | **miR-106b-5p, miR-130b-3p** | **MAPK1** |
| **17** | **miR-106b-5p, miR-376a-5p** | **PTEN** |
| **18** | **miR-93-5p, miR-151a-3p** | **YWHAQ** |
| **19** | **miR-151-a-3p, miR-151b** | **AKT2** |
| **20** | **miR-320c, miR-320d** | **RPS6KA1** |

**Table 2.** From KEGG pathway enrichment analysis mRNAs are predicted. mRNAs that are regulated by more than one shortlisted miRNAs for senescence genes are given in the table.

|  | **miRNAs** | **Predicted Targets**<br>**(mRNAs)** |
| --- | --- | --- |
| **1** | **miR-15b-5p, miR-320e, miR-424-5p** | **CCNT1** |
| **2** | **miR-93-5p, miR-106b-5p** | **CD28** |
| **3** | **miR-15b-5p, miR-221-3p** | **FOXO1, FOXO3** |
| **4** | **miR-31-3p, miR-93-5p** | **FRS2** |
| **5** | **miR-93-5p, miR-107** | **JAK1** |
| **6** | **miR-106b-5p, miR-130b-3p, miR-93-5p, miR-106b-5p** | **MAPK1, MAPK9** |
| **7** | **miR-320c, miR-320d** | **RPS6KA1** |
| **8** | **miR-93-5p, miR-106b-5p** | **RPS6KA5** |
| **9** | **miR-93-5p, miR-106b-5p, miR-130b-3p** | **TGFBR2** |

**Figure 3.**
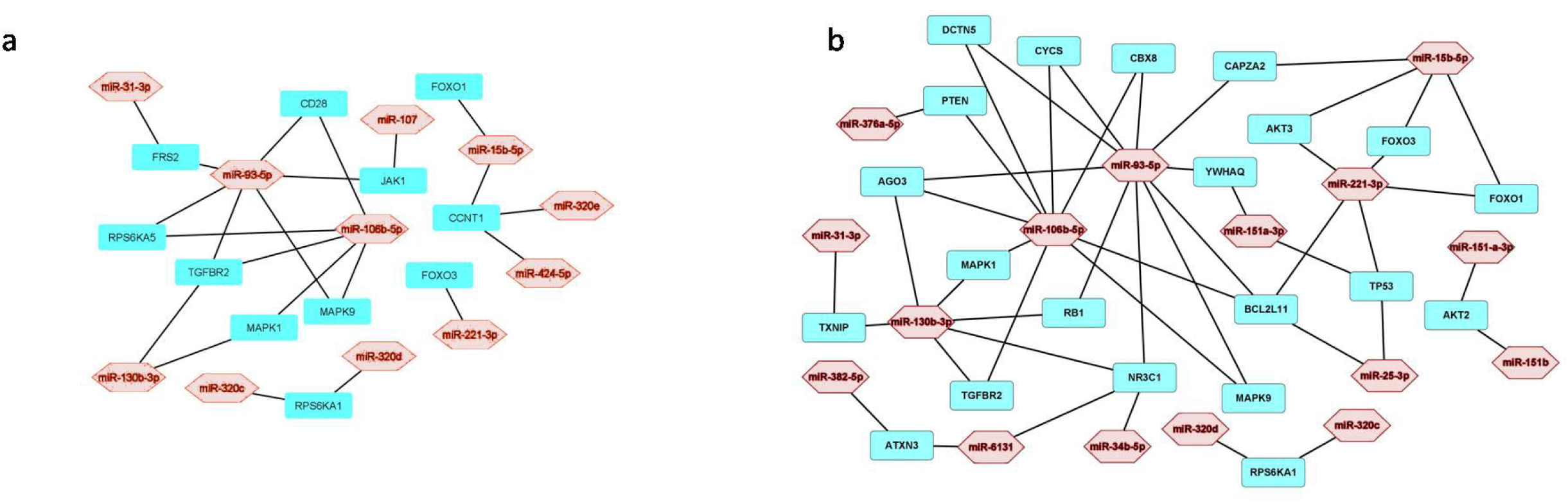
Network image show the relationship between the miRNAs and target mRNAs that are downregulated on TAT+AEA treatment. Network image of (a) Inflammasome related miRNAs - mRNAs and (b) senescence related miRNAs-mRNAs.

### Pathway enrichment analysis

Genes predicted by miRWalk were taken for the KEGG pathway enrichment analysis (Figure 4a-c). Significant pathways predicted by KEGG are related to cell cycle, various cancer pathways, TNF pathway, cellular senescence, viral infection, PI3K-Akt pathway, Human immunodeficiency virus 1 infection, TGF beta pathway.

**Figure 4.**
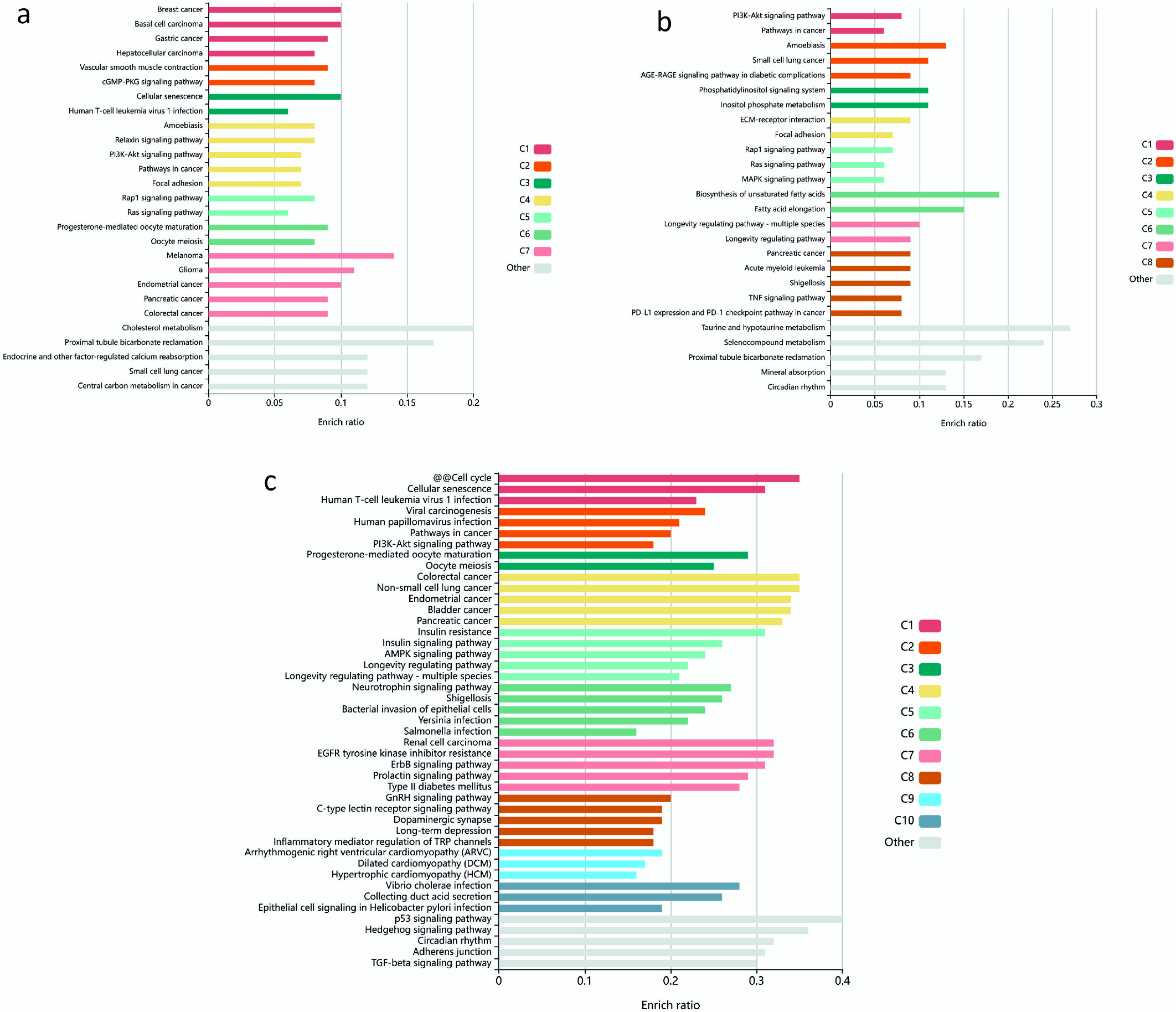
(a) KEGG pathway enrichment analysis was done for the mRNAs that are predicted as target for 122 miRNAs upregulated on TAT treatment. (b) KEGG pathway enrichment analysis was done for the mRNAs that are predicted as targets for 57 miRNAs downregulated with TAT+AEA. (c) KEGG pathway enrichment analysis was done for the mRNAs that are predicted as target for 37 downregulated miRNAs which are common to TAT and TAT+AEA. In the enrichment graph, each row represents enriched function, and the length of the bar represents the enriched ratio. Colored bars represent different clusters.

### Protein-protein interaction (PPI) network analysis

We constructed a PPI network by an online search tool to detect the interacting genes/proteins (STRING) (https://www.string-db.org/). According to PPI network, interactions between inflammatory proteins translated were identified from mRNAs in the miRNA network. FOXO1, FOXO3, MAPK1, MAPK9, JAK1 FRS2, RPS6KA5 and RPS6KA1 are connected with each other significantly (Figure 5a). STRING image shows MAPK1 acts as a junction protein that connects other proteins shortlisted. This is also borne out by the authors’ earlier experimental work in Muller glia at the retina (Krishnan et al. 2012).

**Figure 5.**
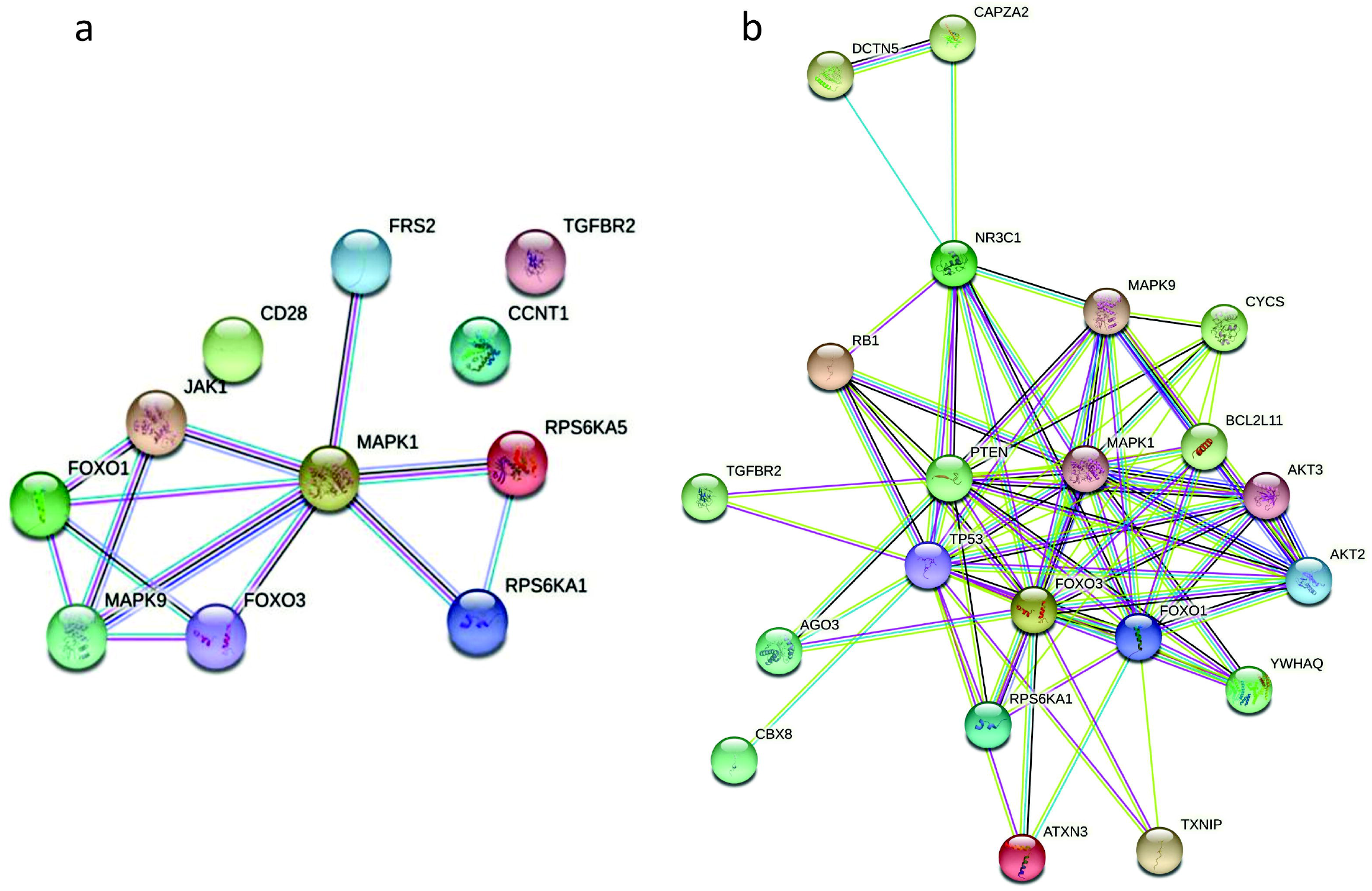
STRING image show protein-protein interactions for the target genes that are regulated by more than one shortlisted miRNAs. STRING image (a) Inflammasome related genes (b) Senescence related genes.

Interactions between senescence related mRNAs using PPI network identified FOXO1, FOXO3, NR3C1, ATXN3, CAPZA2, AKT3, BCL2L11, TP53, TXNIP, CYCS, MAPK9, AGO3, DCTN5, RB1, CBX8, TGFBR2, MAPK1, PTEN, YWHAQ, AKT2 and RPS6KA1 (Figure 5b).

### Comparative analysis of the target miRNAs

Our study was designed to examine miRNAs and their target genes on treatment with endocannabinoid AEA in TAT stimulated human astrocytes. As a part of comparative analysis of our micro array analysis, we predicted the target miRNAs for the mRNA components involved in inflammasome (GSDMD, NLRP1, CARD8, IL-1α, IL1-β, IL-18, AIM2 and, NLRX1), senescence (SIRT1, FOXO3, FOXO4 and CDKN2A) and endocannabinoid system (FAAH, NAPEPLD, MGLL and CNR1). *In silico* prediction of miRNAs by miRDB for these signaling pathway components showed commonality in sets of miRNAs which are present in our experimentally obtained astrocyte data. miRNAs that are common in miRDB and in TAT, TAT+AEA are listed in the Supplementary Table 2. miRNAs that are common in our microarray and miRDB of Inflammasome are hsa-miR-106b-5p, hsa-miR-93-5p, hsa-miR-495-3p, hsa-miR-4306, hsa-miR-15b-5p and hsa-miR-424-5p. Common miRNAs of endocannabinoid system are hsa-miR-106b-5p, hsa-miR-151a-3p, hsa-miR-15b-5p, hsa-miR-34b-5p, hsa-miR-424-5p and, hsa-miR-93-5p. Common miRNAs of senescence are hsa-miR-154-3p, hsa-miR-6131, hsa-miR-361-3p, hsa-miR-4306, hsa-miR-495-3p and hsa-miR-590-5p. This comparative analysis confirm that the miRNAs that are differentially regulated in TAT and TAT+AEA treatment in our microarray study are really related to inflammasome, senescence and endocannabinoid system.

## Discussion

Dysregulated miRNAs contribute to chronic inflammation in the brain, thereby leading to progression of neurological diseases. Reactive glial populations lead to amplified and prolonged neuroinflammatory response (Henry et al. 2008, Huang et al. 2009). Our current study show microRNA involvement in inflammation and senescence phenotype change on endogenous cannabinoid addition in TAT treated astrocytes. Engagement of anti-inflammatory factors after addition of AEA is mediated by multiple microRNAs. Several studies have confirmed that endogenous cannabinoids can reduce inflammation in glial cells through controlling signalling machinery at multiple levels of oxidative stress. Anti-inflammatory action of AEA and their agonists such as WIN-55212-2 can mediate by suppressing pro-inflammatory factors like Interleukin 1 Beta (Cambronero et al. 1989) and expressing IL-6 (Fields et al. 2022). To date, numerous miRNAs have been shown to be significantly up- or down-regulated during aging; many of these miRNAs such as miR-71, miRNA lin-4 were first identified in *C. elegans* as critical for maintaining lifespan (Boehm et al. 2005). Similarly, miR-17-92 in mammals have been shown to be regulators of aging and cellular senescence in various co-morbidities (De Lencastre et al. 2010, Grillari et al. 2010].

Earlier work from our group has already shown the seminal role of MAPK-NF -B axis in AEA and 2-AG alleviating inflammation in Muller glia (Krishnan et al. 2012). Anandamide also reverses barrier properties of Muller cells by controlling production of nitric oxide production (Krishnan et al. 2015). The neuroprotective mechanism involved suppression in production of pro-inflammatory and increase of anti-inflammatory cytokines, mainly through the MAPK pathway (Krishnan et al. 2014). Among the downstream mediators of PI3K/Akt, forkhead box O (FOXO) transcription factors are the main downstream mediators of Akt (Tzivion et al. 2011). FOXOs are negatively regulated by Akt signaling and are known to exert inhibitory effects on cell proliferation in various cell types. Phosphatidylinositol 3-kinase (PI3K), which responds to growth factors and cytokines, has been known to regulate FOXO function. Phosphorylation is the most critical modification as it essentially regulates the translocation of all FOXO proteins between the nucleus and cytoplasm. FOXO translocation to the nucleus and binding to promoter regions of genes that have FOXO response elements is stimulated by the MAP kinase pathway and inhibited by the PI3 kinase/AKT pathway. In the search for regulatory transcription factors that have activity during oxidative stress, FOXO6 has been recently reported for its protective role by inducing antioxidant gene expression during intrinsic and extrinsic aging following IL-10 exposure. Taken together with our previous data in retinal glia this suggests AEA suppresses inflammation by triggering several microRNAs that target proinflammatory cytokine pathways. Our data suggest that dynamic regulatory role of analysed miRNAs serves as a potential mechanistic link between causal involvement of these miRNAs in activation, function and aging.

## Conclusion

In our model we study exacerbation in astrocytes by HIV1 TAT (Sofroniew et al. 2015). To the best of our knowledge this is the first report of AEA -mediated alterations in the miRNAs of activated astroglia. The involvement of miRNAs and their targetome in the process of astrocyte activation, function and the changes affected by AEA is just beginning to be understood. Given that HIV associated neurodegeneration is not limited to neurons, the action of endogenous cannabinoid on astrocytes showing miRNA regulatory role on phenotypes associated with cellular senescence identify alterations in cellular metabolism, secreted cytokines, epigenetic regulation and protein expression suggest important clues to modulating immune response in the CNS.

## Supplementary information

The microarray data discussed in this manuscript has been deposited in the NCBI Gene Expression Omnibus (GEO) under the GEO series accession number GSE210095.

## Acknowledgements

This study was supported by the Department of Biotechnology, Government of India, BT/PR8585/AGR/36/780/2013. DD and KK are supported by the Council of Scientific and Industrial Research.

## Author contributions

Conceived and designed the experiments: DD, NC. Performed the experiments: DD, KK. Analyzed the data: DD, NC, HN. Contributed reagents/materials/analysis tools: SSN, NC. Wrote the paper: DD, HN, NC. The authors have no conflicting interests.

## Abbreviations

miRNAs: MicroRNAs
EC: Endocannabinoid
AEA: N-arachidonoylethanolamine
CNS: Central nervous system
SIV: Simian immunodefeciency virus
HIV: Human immuno deficiency virus
ART: Anti retroviral therapy
TAT: Trans-activator of transcription protein
2-AG: 2-Arachidonoylglycerol
KEGG: Kyoto Encyclopedia of Genes and Genomes
KOBAS: KEGG Orthology-Based Annotation System
STRING: The Search Tool for the Retrieval of Interacting Genes miRDB- MicroRNA Target Prediction Database
HAND: HIV-associated neurocognitive disorder
UTR: Untranslated region

## LEGENDS

Supplementary Table 1 Number of mRNAs targeted by the shortlisted miRNAs based on their Coding DNAs, 3’UTR and 5’UTR are predicted by using miRWAlk.

Supplementary Table 2 In comparative analysis of our microarray analysis, we predicted the target miRNAs for the mRNA components involved in inflammasome (GSDMD, NLRP1, CARD8, IL-1α, IL1-β, IL-18, AIM2 and, NLRX1), senescence (SIRT1, FOXO3, FOXO4 and CDKN2A) and endocannabinoid system (FAAH, NAPEPLD, MGLL and CNR1) by using miRDB. *In silico* prediction of miRNAs by miRDB for these signaling pathway components showed commonality in sets of miRNAs which are present in our experimentally obtained astrocyte data. miRNAs that are common in miRDB and in TAT, TAT+AEA are listed in the Table.

