## Supplementary figures and images for "Antagonistic activity of AEA on TAT treated Human Astrocytes identify Inflammaging pathways"

### Supplemental Table 1

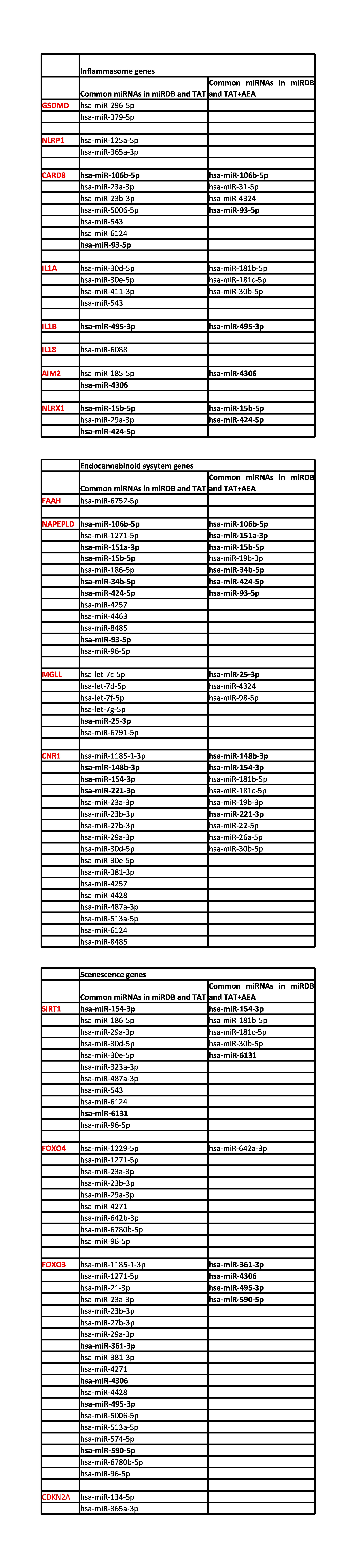

### Supplemental Table 2

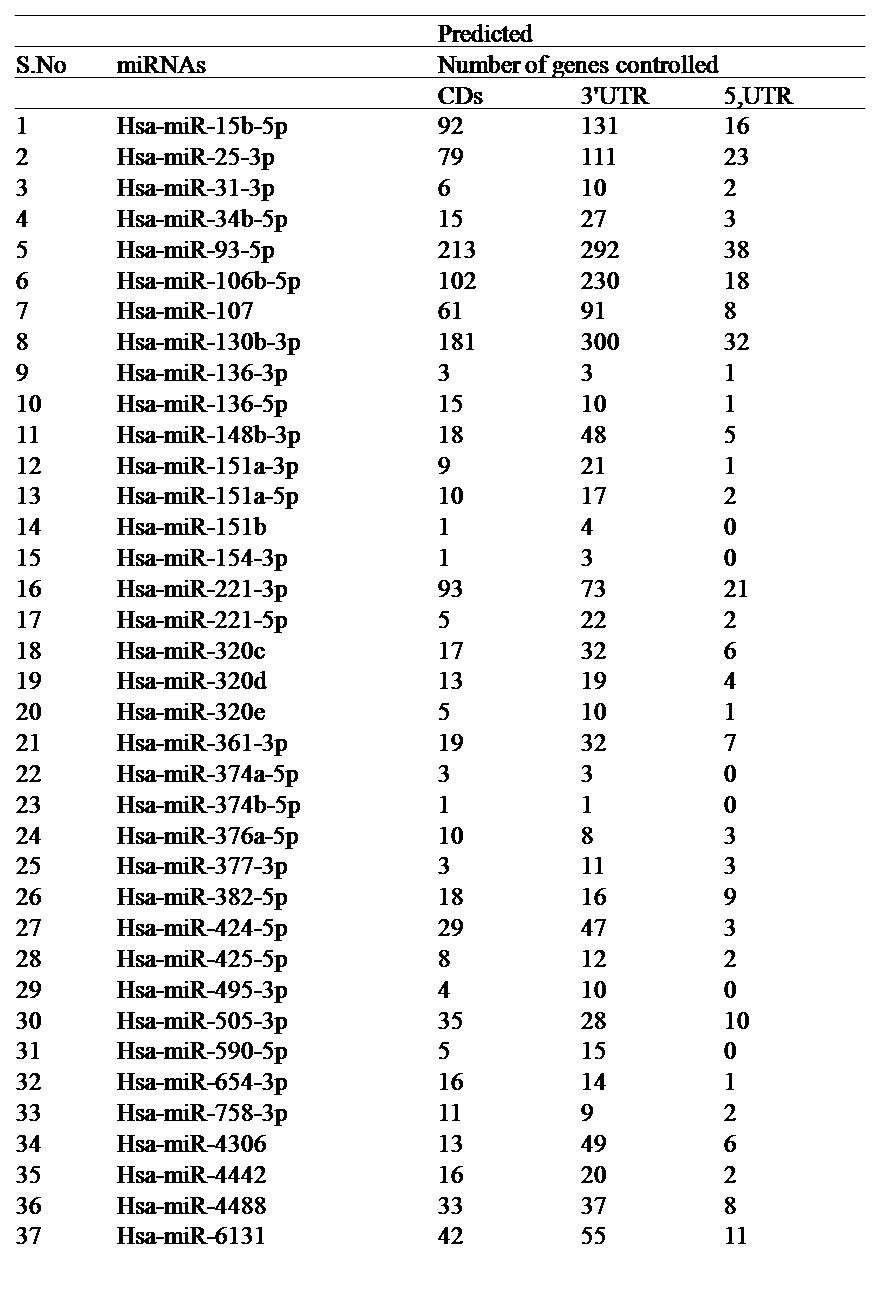
